# Brain connectivity during Alzheimer’s disease progression and its cognitive impact in a transgenic rat model

**DOI:** 10.1101/690180

**Authors:** Emma Muñoz-Moreno, Raúl Tudela, Xavier López-Gil, Guadalupe Soria

## Abstract

The research of Alzheimer’s disease (AD) in their early stages and its progression till symptomatic onset is essential to understand the pathology and investigate new treatments. Animal models provide a helpful approach to this research, since they allow for controlled follow-up during the disease evolution. In this work, transgenic TgF344-AD rats were longitudinally evaluated starting at 6 months of age. Every 3 months, cognitive abilities were assessed by a memory-related task and magnetic resonance imaging (MRI) was acquired. Structural and functional brain networks were estimated and characterized by graph metrics to identify differences between the groups in connectivity, its evolution with age, and its influence on cognition. Structural networks of transgenic animals were altered since the earliest stage. Likewise, aging significantly affected network metrics in TgF344-AD, but not in the control group. In addition, while the structural brain network influenced cognitive outcome in transgenic animals, functional network impacted how control subjects performed. TgF344-AD brain network alterations were present from very early stages, difficult to identify in clinical research. Likewise, the characterization of aging in these animals, involving structural network reorganization and its effects on cognition, opens a window to evaluate new treatments for the disease.

**AUTHOR SUMMARY:** We have applied magnetic resonance image based connectomics to characterize TgF344-AD rats, a transgenic model of Alzheimer’s disease (AD). This represents a highly translational approach, what is essential to investigate potential treatments. TgF344-AD animals were evaluated from early to advanced ages to describe alterations in brain connectivity and how brain networks are affected by age. Results showed that aging had a bigger impact in the structural connectivity of the TgF344-AD than in control animals, and that changes in the structural network, already observed at early ages, significantly influenced cognitive outcome of transgenic animals. Alterations in connectivity were similar to the described in AD human studies, and complement them providing insights into earlier stages and a plot of AD effects throughout the whole life span.

## INTRODUCTION

Alzheimer’s disease (AD) is a neurodegenerative disease related to most cases of dementia in elderly population. Brain damage associated to AD starts decades before the symptomatic onset and clinical diagnose, which has led to consider AD as a continuum (Dubois et al., 2016; Jack et al., 2018). This fact makes the research of disease progression from very early stages a key point to understand AD and develop potential pharmacological treatments or other type of interventions. Consequently, it is essential the identification before the diagnose stage of cohorts of at-risk population to follow them up during years until AD symptoms appear. Recently, studies performed on AD risk population such as carriers of apolipoprotein E (APOE)-*ε*4 or rs405509 alleles have detected brain differences between these subjects and control subjects in elderly (Chen et al., 2015; Reiter et al., 2017; Shu et al., 2015) and middle age population (Cacciaglia et al., 2018; Habib et al., 2017; Mak et al., 2017; ten Kate et al., 2016). However, the identification of at-risk cohorts is challenging and the time required to follow-up them from middle-age to the eventual advanced phase of AD hinders the characterization of the disease progression in patient cohorts. In this sense, animal models provide a helpful approach to evaluate the development of AD from early to advanced stages (Do Carmo & Cuello, 2013; Galeano et al., 2014; Leon et al., 2010; Sabbagh, Kinney, & Cummings, 2013). These models allow the study of earlier stages of the disease, as well as the follow-up of the same subjects during the whole extent of the disease in a relatively short period. An example of AD animal model are TgF344-AD rats. They progressively manifest most pathological hallmarks of the disease including amyloid plaques, tau pathology, oligomeric amyloid *β* (A*β*), neuronal loss and behavioral impairment (Cohen et al., 2013; Drummond & Wisniewski, 2017). Therefore, it is a very suitable model to evaluate AD progression during aging.

Along with the choice of proper animal models, the use of replicable techniques in experimental and clinical research can improve translationality (Sabbagh et al., 2013). This is a crucial point given the high failure rates in the translation between preclinical and clinical trials reported in drug research for AD (Drummond & Wisniewski, 2017; Windisch, 2014). Neuroimaging techniques have been extensively used to identify alterations associated to the disease in a non-invasive way and can be applied in both animal and human cohorts (Sabbagh et al., 2013). In the search for AD biomarkers, magnetic resonance imaging (MRI) has represented a helpful technique to characterize in-vivo brain changes during AD progression (Jack et al., 2015), such as atrophy (Frisoni, Fox, Jack, Scheltens, & Thompson, 2010) or tissue changes (Weston, Simpson, Ryan, Ourselin, & Fox, 2015). In addition, MRI can be used to identify and describe structural and functional brain networks. For this reason it has been used to investigate and support the hypothesis of AD as a disconnection syndrome (Brier et al., 2014; Gomez-Ramirez & Wu, 2014; Palesi et al., 2016; Xie & He, 2012), suggesting that cognitive decline in AD is related to functional or structural disconnection between regions rather than localized changes in specific isolated brain areas. Thus, impairment in structural connectivity associated to AD has been described based on gray matter patterns evaluated in structural MRI or, mainly, based on the fiber tract estimations obtained from diffusion weighted MRI (Daianu et al., 2013; Fischer, Wolf, Scheurich, & Fellgiebel, 2015; Lo et al., 2010; Wee et al., 2011). Likewise, functional disconnection has been evaluated using resting-state functional MRI (rs-fMRI) and graph theory to estimate and quantify functional network (Brier et al., 2014; Sanz-Arigita et al., 2010; Supekar, Menon, Rubin, Musen, & Greicius, 2008); identifying resting-state networks using independent component analysis (Badhwar et al., 2017); or characterizing connectivity between a specific region and the rest of the brain (Gour et al., 2014). Indeed, alterations in structural and functional brain network properties have been described in TgF344-AD animals at early ages (5-6 months) (Muñoz-Moreno, Tudela, Lopez-Gil, & Soria, 2018) as well as differences in functional connectivity of specific regions or networks at several time points from 6 to 18 months (Anckaerts et al., 2019; Tudela, Muñoz-Moreno, Sala-Llonch, Lopez-Gil, & Soria, 2019).

Among the MRI-based studies of AD as a disconnection syndrome, graph theory metrics have become one of the most applied methods to investigate the organization at a global level of both structural (Daianu et al., 2013; Fischer et al., 2015; Lo et al., 2010; Muñoz-Moreno et al., 2018; Wee et al., 2011) and functional (Brier et al., 2014; Muñoz-Moreno et al., 2018; Sanz-Arigita et al., 2010; Supekar et al., 2008) brain networks. Graph metrics provide a quantitative description of different aspects of the network such as integration, segregation or strength, and they allow to perform similar and comparable analyses in structural and functional networks. Since global graph metrics quantify the whole brain structure, they are more sensitive to global reorganization of the networks rather than isolated alterations in individual connections (Rubinov & Sporns, 2010).

Therefore, in the present study, we use graph theory to investigate how the disease progression affects both structural and functional brain networks and its effects in cognitive abilities. In this line, we evaluated how the connectivity impairments observed in young TgF344-AD animals in our previous work (Muñoz-Moreno et al., 2018) evolve during aging. Structural and functional MRI acquisitions were acquired every 3 months from 6 to 18 months of age in a cohort of TgF344-AD and control rats to perform a longitudinal analysis of brain connectivity. Cognitive skills were also evaluated every 3 months to test the impact of connectivity alterations in cognition. Hence, this work aims to contribute to the understanding of AD progression and its association with cognitive decline from the perspective of the disease as a disconnection syndrome.

## MATERIALS AND METHODS

### Subjects

The experiments were performed in a cohort of 18 male Fisher rats including TgF344-AD animals (Cohen et al., 2013) and their wild-type littermates, which were evaluated at 5 time points. Table 1 shows the details on the resulting sample size and average age per time point after MRI experiments.

**Table 1.** Sample size and age (median ± interquartile range) at each of the 5 acquisitions. Sample size in time point 1 is different in the structural and functional analysis (*corresponds to the number of rs-fMRI acquisitions). Variability in the age of acquisition at the first time point is due to differences in the cognitive training duration. Age difference between the groups was not significant.

|  | Control |  | TgF344-AD |  |
| --- | --- | --- | --- | --- |
|  | N | Age (months) | N | Age(months) |
| Time point 1 | 8/5* | 5.33±0.3 | 8/7* | 6.33±1.22 |
| Time point 2 | 9 | 9.2±0.5 | 8 | 8.67±0.2 |
| Time point 3 | 7 | 11.3±0.22 | 9 | 11.3±0.03 |
| Time point 4 | 8 | 14.9±0.425 | 9 | 14.87±0.03 |
| Time point 5 | 8 | 17.92±0.52 | 6 | 18.03±0.62 |

The animals were housed in cages under controlled temperature (21 ± 1°C) and humidity (55 ± 10%) with a 12-hour light/12-hour dark cycle. Food and water were available ad libitum during all the experiment, except during behavioral test periods as explained below. At 2 months of age, animals start a cognitive training to perform delay non-matched sample (DNMS) task. Once the learning criteria was achieved, their performance in DNMS was evaluated and the first MRI scan was acquired. Afterwards, two weeks of DNMS sessions followed by MRI acquisition were repeated every three months resulting in five evaluated time points.

### Cognitive function evaluation

Every three months, working memory was evaluated by DNMS test, following the procedure described in Muñoz-Moreno et al. (2018). DNMS was carried out in isolated operant chambers (Med Associates, USA), equipped with a pellet dispenser and three retractable levers, two of them in the wall where the pellet is (right and left levers) and the other in the opposite side (center lever). During the testing weeks, rats were food-deprived, receiving 75% of the usual food intake. In brief, DNMS requires the animal to press the levers following a specific sequence to obtain a pellet. First, in the sample phase, right or left lever appeared and after the animal pressed it, a delay randomly timed between 1 and 30 seconds started, after which the center lever appeared. When the animal pressed it both right and left levers were extended again. Correct response required a press on the lever opposite to the presented in the sample phase. Each DNMS session finished after 90 minutes or when 90 trials were completed. The number of trials and percentage of correct responses were recorded.

Before the first test, animals underwent a habituation and training phase to acquire the required skills. This phase started when the animals were two months old and finished when they achieved an acquisition criteria as explained in Muñoz-Moreno et al. (2018). After that, animals underwent 15 DNMS sessions (five sessions per week) to evaluate and consolidate the learnt task. At each of the following four time points, animals performed 10 DNMS sessions (two weeks).

### Magnetic Resonance Imaging

MRI acquisitions were performed on a 7.0T BioSpec 70/30 horizontal animal scanner (Bruker BioSpin, Ettlingen, Germany). Animals were placed in the supine position in a Plexiglas holder with a nose cone for administering anesthetic gases (1.5% isoflurane in a mixture of 30% O_2_ and 70% *CO*) and were fixed using tooth and ear bars and adhesive tape. The rat received a 0.5 ml bolus of medetomidine (0.05 mg/kg; s.c.) and a catheter was implanted in its back for continuous perfusion of medetomidine. Isoflurane was gradually decreased until 0% and 15 minutes after the bolus the medetomidine perfusion (0.05 mg/kg; s.c.) started at rate 1 ml/hour. The acquisition protocol included:

▪ T2-weighted images, acquired using a RARE sequence with effective echo time TE = 35.3 ms, repetition time TR = 6000 ms and RARE factor = 8, voxel size = 0.12 × 0.12 mm^2^, 40 slices, slice thickness = 0.8 mm and field of view FoV = 30×30×32 mm^3^.
▪ T1-weighted images, acquired using an MDEFT protocol with TE = 2 ms, TR = 4000 ms, voxel size = 0.14×0.14×0.5 mm^3^ and FoV=35×35× 18 mm^3^.
▪ Diffusion weighted images (DWI) using a spin-echo EPI sequence with TE = 24.86 ms, TR = 15000 ms, four segments, 60 gradient directions with b-value = 1000 s/mm^2^ and five volumes with b-value = 0 s/mm^2^; voxel size = 0.31 ×0.31 ×0.31 mm^3^ and FoV = 22.23×22.23× 18.54 mm^3^.
▪ rs-fMRI using a gradient echo T2* acquisition, with the following parameters: TE = 10.75 ms, TR = 2000 ms, 600 volumes (20 minutes), voxel size = 0.4× 0.4× 0.6 mm^3^, FoV = 25.6 × 25.6 × 20.4 mm^3^. rs-fMRI acquisition is scheduled after anatomical and diffusion MRI to ensure that isoflurane dose has been removed and the animals are sedated only by medetomidine.

To illustrate the acquisition quality, Supplementary Figure 1 shows a series of slices of colored fractional anisotropy computed in a randomly selected subject of the cohort; and Supplementary Figure 2 displays a selection of functional networks extracted from the whole cohort using independent component analysis (ICA).

### Image processing and connectome definition

The acquired images were processed to obtain the structural and functional connectomes following the methodology described in Muñoz-Moreno et al. (2018). Briefly, a rat brain atlas was registered to the T2-weighted images to obtain brain masks and region parcellations (Schwarz et al., 2006). T1-weighted images were used to segment the brain into white matter (WM), gray matter (GM) and cerebrospinal fluid (CSF) based on tissue probability maps registered from an atlas (Valdés-Hernández et al., 2011) to each subject brain. Parcellation and segmentation were registered from T2/T1-weighted volumes to DWI and rs-fMRI spaces to define the regions between which connectivity was assessed.

Fiber tract trajectories were estimated from DWI volumes using deterministic tractography based on constrained spherical deconvolution model, considering WM voxels as seed points. Dipy was used to process DWI volumes and estimate the fiber tracts (Garyfallidis et al., 2014). The resulting number of streamlines generated per subject are of the order of 10^5^ (6.18·10^5^± 0.67·10^5^). As defined in Muñoz-Moreno et al. (2018), the structural connectome included 76 regions. Connection between two regions was defined if at least one streamline started in one region and ended in the other. The resulting structural connectomes have an average density of 62.22±2.78. Three connectomes were considered according to the connection weight definition:

▪ Fractional anisotropy (FA) weighted connectome (FA-w): The connection weight between two regions is defined as the average FA in the streamlines connecting them.
▪ Fiber density (FD) weighted connectome (FD-w): The connection weight between two regions is computed as the number of streamlines normalized by the region volumes and the streamline length (Muñoz-Moreno et al., 2018).
▪ Structural binary connectome: connection weight is 1 between connected regions and zero otherwise.

rs-fMRI was processed to obtain the average time series in the GM voxels of each of the regions of interest. Preprocessing includes slice timing, motion correction by spatial realignment using SPM8, and correction of EPI distortion by elastic registration to the T2-weighted volume using ANTs (Avants, Epstein, Grossman, & Gee, 2008). Afterwards, NiTime (http://nipy.org/nitime/) was used for z-score normalization and detrending of the time series, smoothing with an FWHM of 1.2 mm, frequency filtering between 0.01 and 0.1 Hz, and regression by motion parameters and WM and CSF average signals. Since brain activity identified by rs-fMRI has been constrained to GM (Power, Plitt, Laumann, & Martin, 2017), only regions comprising GM tissue were considered as nodes in the functional connectome (54 regions). Both weighted and binary functional connectomes were defined. The connection weight was the partial correlation between the pair of regional time series, transformed by Fisher’s *z*-transformation. Negative correlation coefficients were excluded since the proposed analysis is based on graph theory metrics that are not defined for signed edges (Fornito, Zalesky, & Breakspear, 2013). All the connections with positive weight (*z* > 0) were considered. Binary functional connectome was defined setting to one connections were *z* > 0 and zero otherwise. The resulting functional connectomes have an average density of 27.72±0.53.

### Brain network analysis

Brain network organization was described using graph theory metrics, including degree, strength, clustering coefficient and local and global efficiency. These metrics provide a description of different aspects of brain networks at a global level: network density, integration and segregation (Rubinov & Sporns, 2010).

Nodal degree measures the number of connections of a region. Nodal strength is computed as the sum of the region connection weights. Network degree and strength are respectively the nodal degree and strength averaged in all the brain regions. Higher network degree/strength indicates more or stronger connections. Global efficiency measures network integration: the ability to combine information from different regions. It is inversely related with the shortest path length between each pair of nodes. Higher global efficiency characterizes stronger and faster communication through the network. Network segregation, the ability for specialized processing within densely interconnected groups of regions, was quantified by local efficiency and clustering coefficient. Local efficiency is the average in the whole network of the nodal efficiencies (efficiency of the subnetwork associated to a region). Clustering coefficient is the average of the nodal clustering coefficients, which measure the number of neighbors of a node that are also neighbors with each other. High values of these metrics are related to highly segregated and connected networks. In this way, alterations in the whole-brain organization were evaluated using the same kind of measures in both structural and functional connectomes. Nonetheless,the interpretation of these parameters must take into account the specific kind of connectome and the definition of the connection weights. On the other hand, these metrics have been commonly used in human studies of AD (Brier et al., 2014; Daianu et al., 2013; Fischer et al., 2015; Sanz-Arigita et al., 2010), and therefore, can provide comparable and translational results.

### Statistics. Longitudinal models

The main objective of our study was the research of alterations in the longitudinal evolution of the TgF344-AD brain networks with respect to the control group and its impact on cognition. Linear mixed effects (LME) models (Oberg & Mahoney, 2007) were used to model the influence of age and group in the network metrics and to identify differences in the effect of aging between the two groups. LME models include both fixed effect (parameters common to an entire population such as age or group) and random effects (subject-specific parameters modeling the subject deviation from the population).

LME was defined to regress each of the network metrics including group, age and the interaction between them as independent variables:

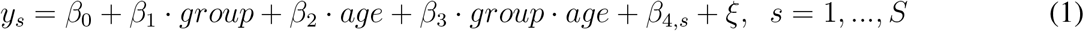

where *y_s_* is the network metric in the subject *s* at a given age, *s* represents each of the *S* subjects, *β*_0_ is the global intercept, *β*_1_, *β*_2_, *β*_3_ the fixed effect parameters, assessing the influence of group, age and interaction respectively, and *β*_4,*s*_ the subject specific correction. is the regression error term.

Multiple comparisons were corrected using false discovery rate (FDR) (Benjamin & Hochberg, 1995). The effects of group, age or interaction between age and group were considered significant if the corrected p-value (*p_FDR_*) was less than 0.05. When the interaction was significant, control and transgenic groups were modeled separately to evaluate the age effect in each group:

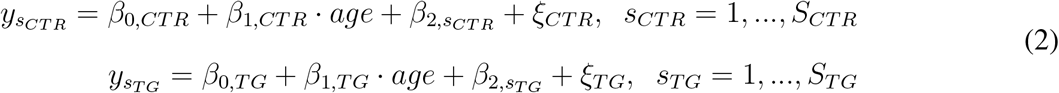

To complement the longitudinal analysis, differences in brain networks at each of the five acquisition time points were evaluated. Kruskal-Wallis tests were applied to identify statistically significant differences between whole-brain organization in transgenic and control groups. Multiple comparisons were corrected using FDR (Benjamin & Hochberg, 1995). Together with this, Network-Based Statistics (NBS) toolbox (Zalesky, Fornito, & Bullmore, 2010) was used to identify specific networks of connections which differ between transgenic and control animals. NBS was performed with the following settings: t-test with a threshold of 3.1, 5000 permutations and a significance level of *p* < 0.05.

Finally, the relationship between connectivity and cognitive performance was evaluated fitting an LME model to test if such relation is significant and if it differs between the groups:

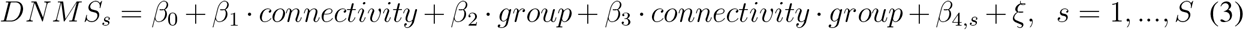

where *DNMS_s_* represents the result in the DNMS task (number of trials and percentage of correct responses) and connectivity refers to each of the network metrics. Thus, the relationship between cognitive outcome and each of the network parameters was evaluated. Multiple comparisons were corrected by FDR (Benjamin & Hochberg, 1995). When interaction was significant (*p_FDR_* < 0.05), the model was fitted separately to transgenic and control groups to test the significance of the relationship between the network metric and the cognitive outcome in each group.

## RESULTS

### Longitudinal analysis

Linear mixed effects (LME) model was fitted to regress each of the network metrics and to evaluate significant effects of age, group or their interaction. Results are shown in Figure 1, where the whole distribution of data is shown (each point represents a time-point / subject), as well as the fitting of the LME model as a function of group and age. If a significant effect of the interaction between age and group was detected, the model was adjusted to each of the groups independently to fit the network metric as a function of age. Significant *p_FDR_* values of each of the terms in the model (group, age and group-age interaction) are displayed in Figure 1. Supplementary Table 1 compiles *p_FDR_* values and effect sizes (Cohen’s *f*^2^) for the parameters of the LME model fitted to the whole cohort, and Supplementary Table 2 shows *p_FDR_* values and effect sizes (Cohen’s *f*^2^, *R*^2^) of the model adjusted to each of the groups.

**Figure 1.**
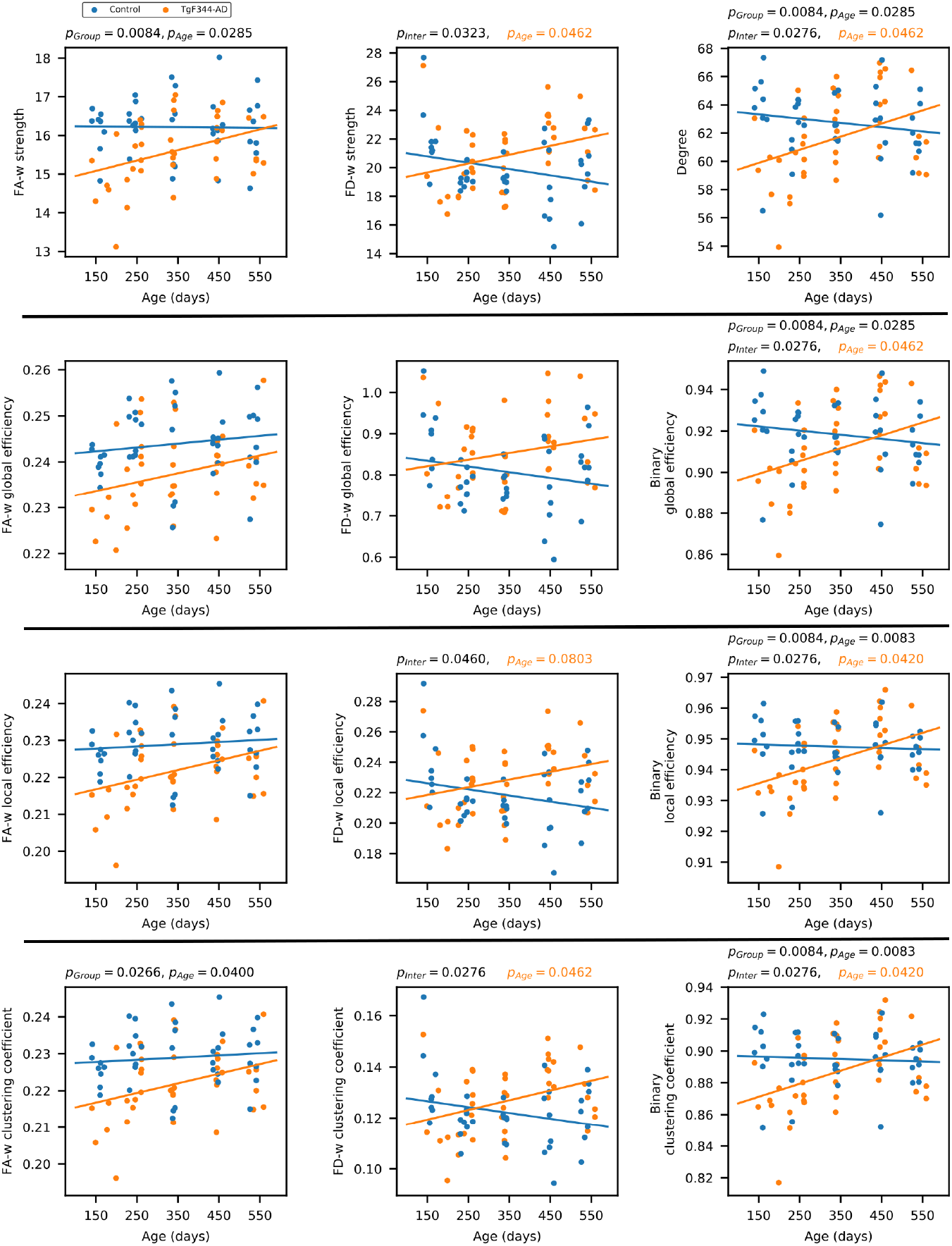
Global network metrics of the structural connectome. Strength, global and local efficiency and average clustering coefficient of the three structural connectomes (fractional anisotropy-weighted, fiber density weighted and binary connectomes). For each network metric, each dot represents the value of one animal at one time point (blue: control, orange: TgF344-AD). Blue and orange lines represents the fit of the linear mixed effects model; *p_FDR_* values are shown for significant effects (*p_group_* group effect, *p_age_* age effect, *p_Inter_* effect of the interaction between age and group). If interaction was significant group models were fitted to the data, in orange, *p_FDR_* values of the effect of age in the LME model fitted to the transgenic group (no significant effects were observed in the control group).

In FA-w connectome, group effect was significant in strength and clustering coefficient, but not significant interaction between group and age was detected. Both FA-w strength and clustering were clearly decreased in the transgenic group. A significant increase of FA-w strength with age was observed, with a more notable slope in the case of transgenic animals.

The interaction between group and age was significant in FD-w strength, local efficiency and clustering coefficient. In all these cases, an increase with age in the transgenic group - significant in strength and clustering coefficient-was detected, opposite to the non-significant decrease observed in controls. Similar behavior was observed in the binary connectome, where significant age-group interaction was observed in all the network metrics, which were significantly affected by age in the transgenic group.

No significant effects of age, group or interaction were detected neither in functional connectivity nor in cognitive performance.

### Group differences by time points

Figure 2 shows the distribution of network metrics between groups at each of the five acquisition time points, and the statistically significant differences. Significance was considered as *p_FDR_* < 0.05. Only structural network metrics are shown, since no differences were found in functional connectomes. Supplementary Table 3 shows the *p_FDR_* values and effect sizes (*η*^2^) of both structural and functional connectomes.

**Figure 2.**
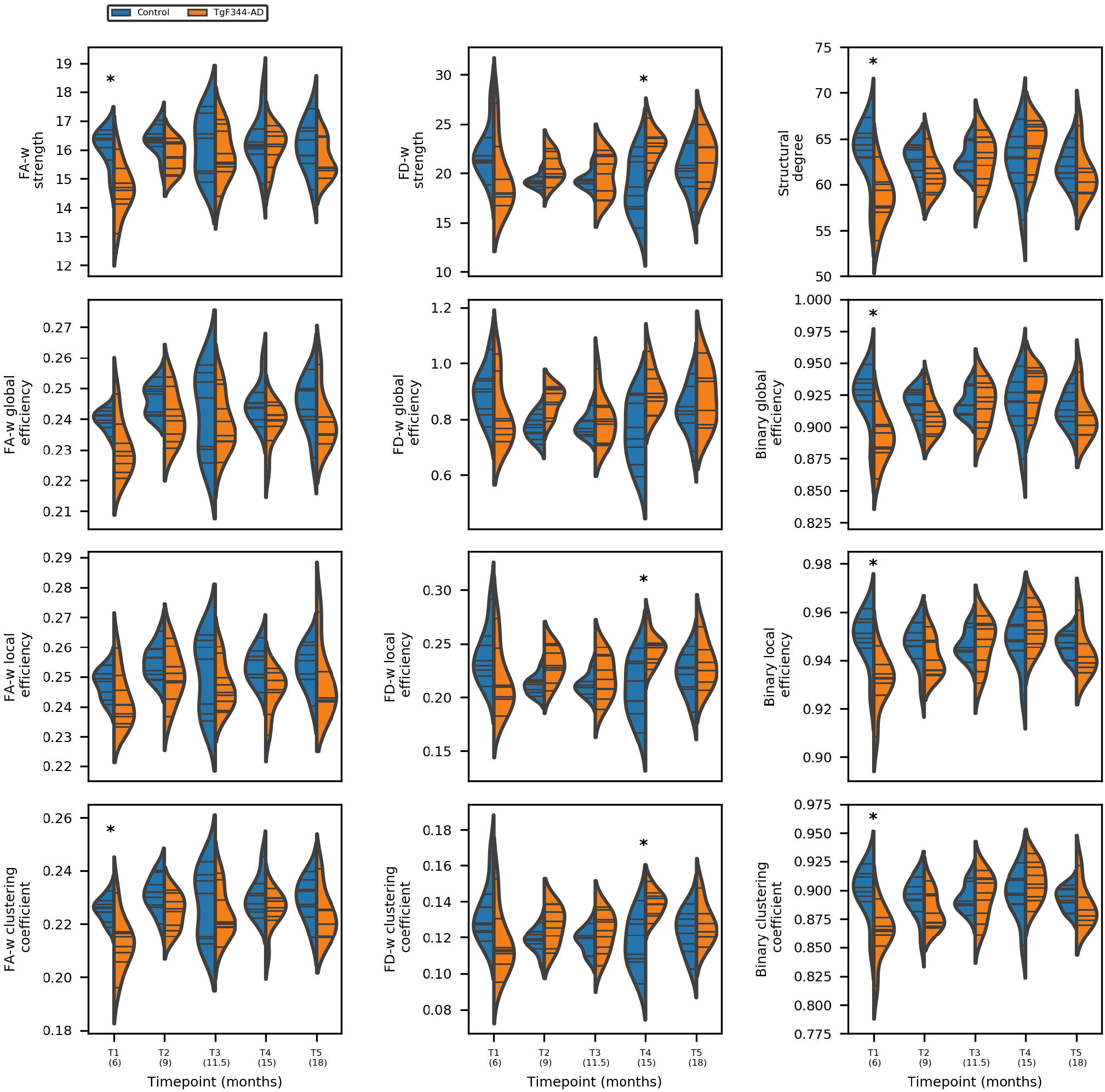
Network metrics of the three structural connectomes (fractional anisotropy weighted, fiber density weighted and binary connectome) in control (blue) and transgenic (orange) groups at each of the five time points. * represents statistically significant difference (*p_FDR_* < 0.05)

Most differences were observed at the earliest time point where FA-w and binary structural network metrics were significantly decreased in the transgenic group with respect to controls. Regarding the FD-w connectome, while a tendency to decreased values were observed in the transgenic group with respect to controls at the first time point, a significant increase was detected in strength, local efficiency and clustering coefficient at 15 months of age.

No significant differences were found in the functional network metrics.

Together with graph metrics we evaluate differences in network edges. Figure 3 shows the group average FA-w and functional weighted connectomes considering only the strongest connections (FA-w> 0.3 and *z* > 0.05, respectively) to provide an illustrative plot of the brain networks. Stronger and denser FA-w connections are observed in the control group in comparison with transgenic, especially in the last time point, although network metrics were not significantly different. A decrease in the connection strength with time in the transgenic brain can also be observed. Regarding the functional connectome it can be seen that at 18 months of age, *z* > 0.05 connections in the TgF344-AD brain are less and weaker. In previous time points connectivity is similar between both groups, and even higher strength in specific links can be observed in the transgenic brain.

**Figure 3.**
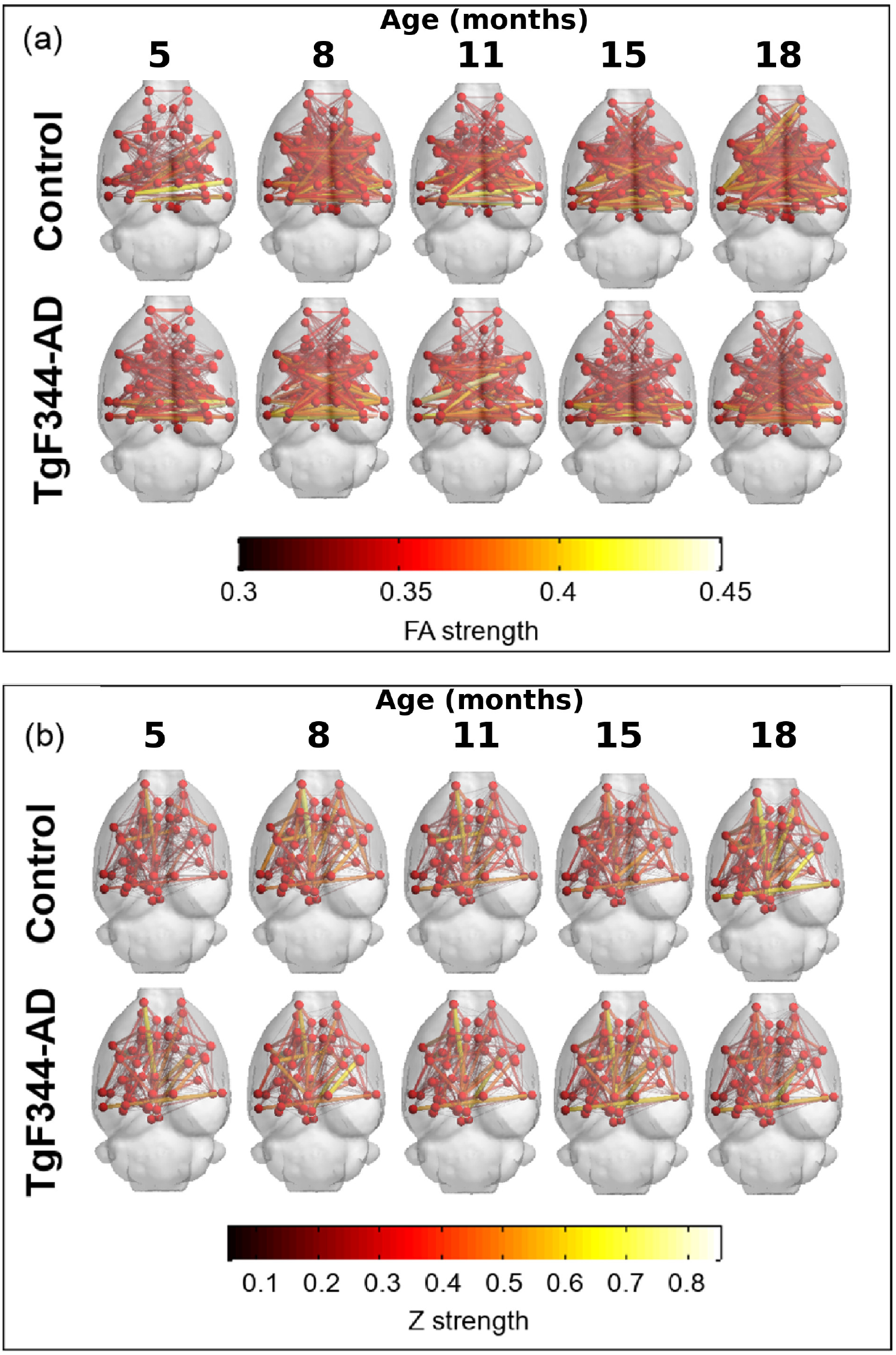
Average structural and functional brain networks. (a) Group average of the FA-w connectome; (b) group average of the functional weighted connectome at each time point. Only the strongest connections are plotted (FA-w> 0.3 and *z* > 0.05). Color and width of the links represent connection strength.

To statistically identify differences in the network edges, NBS was applied. Resulting networks are shown in Figure 4. Differences were detected at 8 months of age (time point 2) in a subnetwork of the FD-w connectome (increased in transgenic animals) and at 15 months of age (time point 4) in FA-w and functional connectomes. While the subnetwork identified in the FA-w connectome was decreased in transgenic animals, subnetworks in the functional network were increased in this group. The list of regions between which connectivity was altered is shown in Supplementary Material.

**Figure 4.**
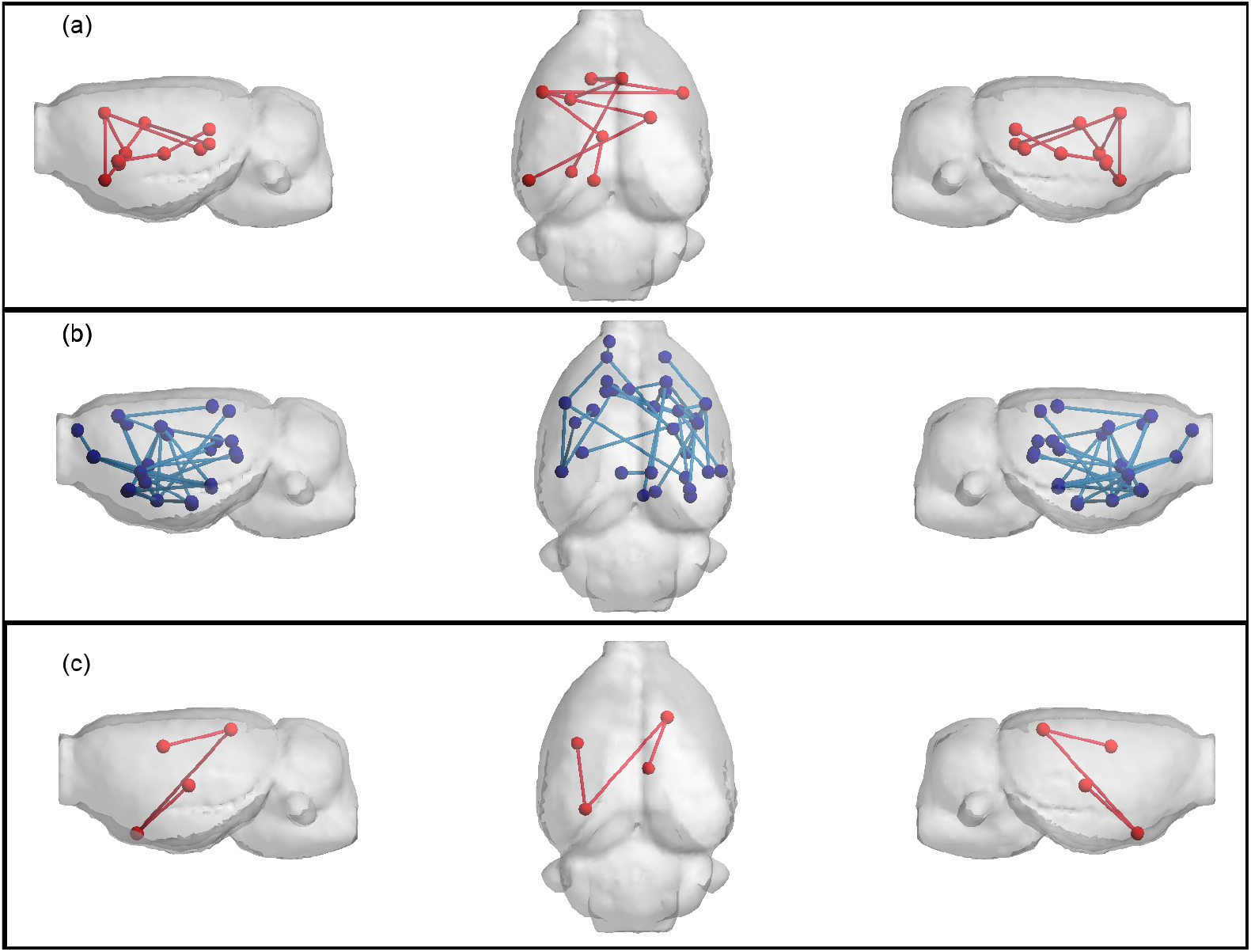
Networks altered in TgF344-AD animals resulting from NBS analysis. a) FD-w connectome at 8 months of age, altered connections are related to limbic system; b) FA-w connectome at 15 months of age, connections related to memory processing and executive functions; c) functional connectome at 15 months of age, connections related to processing of sensory inputs. Red indicates subnetwork increased in the transgenic group; blue, subnetwork decreased in the transgenic group.

### Relationship between connectivity and cognition

The effect of connectivity in cognitive results and if this effect was different in transgenic and control animals was evaluated using LME models.

Although no significant differences were observed in cognitive outcome between groups (see Supplementary Figure 3), it can be noted that: a) transgenic animals performed lower number of trials than control animals at the first time point and b) there is a high variability in the results of the transgenic group at the last time point, when the performance of some of the animals sharply falls. Besides, brain network organization had a significant impact on the cognitive results.

The interaction between FA-w strength and group had a significant effect in DNMS results (*p_inter_* = 0.0223). Considering the group specific model, this metric had a significant influence in the number of trials performed by the transgenic animals (*p_tg_* = 0.0018), but not in controls. Group and metric shown also a significant effect in cognitive outcome (*p_group_* = 0.0203 and *p_metric_* = 0.0006, respectively).

Similar results were observed considering any of the FD-w metrics (strength: *p_group_* = 0.0098, *p_metric_* = 0.0007, *p*_inter_ = 0.0016; local efficiency: *p_group_* =0.0098, *p_metric_* =0.0005, *P_inter_* = 0.0167; clustering coefficient: *p_grous_* = 0.0051, *p_metric_* = 1.97 · 10^-5^, *p_inter_* = 0.0065) or binary metrics (degree: *p_group_* =0.0051, *p_metric_* = 1.6·10^-5^, *p_inter_* =0.0065; global efficiency: *p_group_* =0.0051, *p_metric_* = 1.6 · 10^-5^, *p_inter_* = 0.0065; local efficiency: *p_group_* = 0.0098; *p_metric_* = 4.4 · 10^-6^, *P_inter_* = 0.0064; clustering coefficient: *p_group_* = 0.0098, *p_metric_* = 4.4 · 10^-6^, *p_inter_* = 0.0064), except for FD-w global efficiency (*p_group_* = 0.0051, *p_metric_* = 0.0035), where the interaction between group and metric was not significant. In all the cases, the higher the FD-w or structural binary network metrics, the more trials the transgenic animal performed (FD-w strength, *p_tg_* = 0.0019; local efficiency, *p_tg_* = 0.0014; clustering coefficient, *p_tg_* = 6.54 · 10^-5^ and binary degree, *p_tg_* = 5.29 · 10^-5^; global efficiency, *p_tg_* = 5.29 · 10^-5^; local efficiency, *p_tg_* = 1.26 · 10^-5^; clustering coefficient *p_tg_* = 1.26 · 10^-5^). All the reported p-values are FDR corrected. Figure 5 shows FA-w, FD-w strength and structural degree (results are similar in all the mentioned metrics). More details are provided in Supplementary Tables 4 and 5.

**Figure 5.**
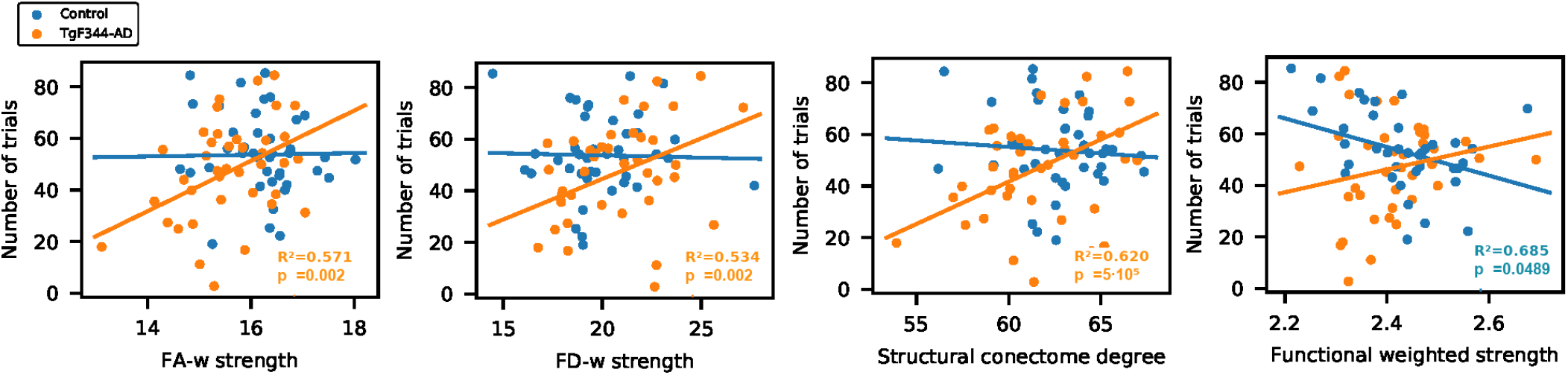
Cognitive performance and network metrics. Linear mixed model fit of DNMS result as a function of network metric and group. Relationship between the structural and functional connectome metrics and number of trials performed in DNMS test. R-squared and *p_FDR_* in case of significant metric effect (orange: significance in TgF344-AD group; blue: significance in control group).

As shown in Figure 5, the number of trials was significantly influenced by the interaction between group and functional weighted strength (*p_inter_* = 0.0150), global efficiency (*p_inter_* = 0.0180) and local efficiency (*p_inter_* = 0.0167). When specific group models were considered, the effect of functional strength in the cognitive performance was significant in the control animals (*p_ctr_* = 0.0489), but not in the transgenic cohort.

## DISCUSSION

There is a growing interest in the study of early stages of AD and its progression until symptomatic onset, since brain changes start decades before the clinical diagnosis (Dubois et al., 2016; Jack et al., 2018). To contribute to the understanding of these early brain changes and their progression during aging, the present study focuses on an animal model of the disease and describes the longitudinal evolution of structural and functional brain network organization from very early stages. Although recent studies have evaluated connectivity in population at risk of AD or in its preclinical phases in human cohorts (Berlot, Metzler-Baddeley, Ikram, Jones, & O'Sullivan, 2016; Farrar et al., 2017; Pereira et al., 2017), they focused on elderly or middle-aged subjects and in case of longitudinal analysis only short periods of time with respect to human life span have been evaluated. In the present study, the use of TgF344-AD rats allows the investigation of earlier alterations and to follow-up subjects during all their life span. Therefore, it can provide new insights into the disease progression and be helpful in the investigation of treatments and interventions.

Our results show differences between transgenic and control groups in the progression of the structural connectivity during aging. Alterations in the structural brain network of population at risk of AD or in its preclinical phases in human cohorts of elderly or middle-aged subjects have been reported (Berlot et al., 2016; Chen et al., 2015; Farrar et al., 2017; Fischer et al., 2015; Pereira et al., 2017; Shu et al., 2015; Zhao et al., 2017), which are in line with the differences we have observed in TgF344-AD animals at equivalent ages (15 months). Furthermore, earlier alterations were observed in the animal model: structural connectivity differences were already present in young animals (six months of age). Together with this, differences in the evolution of the structural metrics were detected. While aging had no significant effect in the evolution of network metrics in the control group, it significantly affected metrics in the transgenic animals. In this group, network metrics increased linearly with age, but a decrease in the metric values at the last time point can be observed in Figure 1 and 2, although LME could not fit this change of trend. In spite of this global increment with age, FA-w network metrics in transgenic animals remain always lower than in controls, in line with which has been observed in preclinical or clinical phases of AD in aged patients (Chen et al., 2015; Pereira et al., 2017; Shu et al., 2015). Binary and FD-w metrics also increased significantly in transgenic animals during aging, but they were decreased with respect to controls only at early ages. Indeed, FD-w was increased in transgenic animals at 8 and 15 months. This could be related with the hyperconnectivity effect described after brain injury and in preclinical phases of AD (Hillary & Grafman, 2017), explained as a mechanism to preserve communication in the network and minimize the behavioral deficits. Although hyperconnectivity has been mainly identified in functional networks, higher binary structural network properties were also described in APOE *ε*4 carriers before mild cognitive impairment (MCI) appears (Ma et al., 2017). Thus, to compensate the lower FA-w strength in the network, more connections would be established to preserve behavioral performance. In this line, the increase in FA-w network metrics with age observed in transgenic animals is probably related to the presence of more connections rather than to FA increase in the brain. The results obtained with NBS also points to this line. They showed hyperconnectivity in subnetworks of functional and FD-w connectomes, while decreased connectivity was observed in a subnetwork of the FA-w connectome. Namely, hyperconnectivity was detected in networks related with the processing of sensory inputs in both functional and FD-w connectome. The altered connections observed in the FD-w network are part of the limbic system and are responsible for the processing of different sensory inputs required to link emotions and memories. This could try to compensate the decrease observed in FA-w connectome in networks related to memory processing and executive functions.

Impairments in functional connectivity have been described in at risk, preclinical or clinical AD cohorts, but depending on methodological issues or cohort selection differences have been reported in opposed senses (Phillips, McGlaughlin, Ruth, Jager, & Soldan, 2015). Likewise, in the studied animal model, TgF344-AD, differences in connectivity between specific regions or networks have been reported (Anckaerts et al., 2019; Tudela et al., 2019) at several ages. In the present work, we focused on the analysis of brain network globally instead of evaluation of specific connections. From this point of view, the whole-brain network metrics describing integration and segregation of the functional connectomes did not significantly differ between control and transgenic groups. This could be related to three factors. First, the hyperconnectivity effect previously mentioned (Hillary & Grafman, 2017) between specific regions or networks to compensate damaged connections (hypo-connectivity). This effect has been described, for instance, in young APOE ε4 carriers (Ma et al., 2017) or amnestic MCI patients (Kim et al., 2015). Since global network metrics involve averaging properties of all the connections (Rubinov & Sporns, 2010), decreased connections could be compensated by increased connections (such as the detected by NBS at 15 months of age), resulting in similar values of the global network metrics, as observed in our study. Second, the training and the repetition of the cognitive task could lead to a learning effect, increasing the cognitive reserve of these animals and therefore preserving the functional connectivity. Higher functional connectivity has been related with higher cognitive reserve in both healthy elders (Arenaza-Urquijo et al., 2013) and MCI patients (Franzmeier et al., 2017) with respect to subjects with low cognitive reserve. This fact could also be related with the absence of significant differences in the cognitive outcome. Nevertheless, further investigation on animals not undergoing DNMS should be performed to confirm this hypothesis. Finally, the third factor that could influence the absence of functional connectivity differences could be the disease timing. Brain changes as amyloid-β concentration and neural loss in TgF344-AD have been described from 16 months of age in Cohen et al. (2013) and significant cognitive impairment is mainly described from 15 months (Cohen et al., 2013; Tsai et al., 2014) although tendencies to impairment or differences in anxiety or learning abilities have been reported at earlier stages (Cohen et al., 2013; Muñoz-Moreno et al., 2018; Pentkowski et al., 2018). Actually, an increase in variability of the cognitive outcome was observed at 18 months of age, when some of the transgenic animals performed much worse than at previous ages. This suggests that the symptomatic onset occurs around 18 months of age, and before this age, functional connectivity might compensate the structural network damage resulting in similar functional network metrics than in control animals. The observed significant relation between structural network metrics and cognitive performance could be related with the ability of functional connectivity to cope with the structural disconnection preserving cognition, until a breakpoint when it is not able to deal with extensive structural network damage. This relationship between structural connectivity and cognition has been also described in MCI subjects (Berlot et al., 2016; Farrar et al., 2017) and in cognitive normal individuals harboring amyloid pathology and neurodegeneration (Pereira et al., 2017), while no correlation was observed in control subjects. Coherently, structural connectivity did not correlate with DNMS in our control cohort, where correlation between DNMS and functional connectivity was observed.

### Strengths and limitations

The use of MRI-based connectomics to longitudinally analyse the brain network in an animal model of AD provides a valuable and highly translational approach to the research on the mechanism of the disease progression, since the applied methodology can be easily translated to clinical investigation. Besides, animal models allows for characterization and follow-up of the same cohort from early stages until advanced phases of the disease. Indeed, the model used in our experiments, the Tg344-AD rats, has been shown to develop all the AD pathological hallmarks in a progressive manner, which makes it especially suitable for longitudinal evaluation.

The use of graph metrics allows for comparison between alterations observed in the animal model and previous results in human cohorts in literature. This is essential to validate how the animal model mimics the pathology in patients. It is also a critical point to allow translationality between preclinical and clinical trials in the research for AD treatments (Drummond & Wisniewski, 2017; Sabbagh et al., 2013). Our results are coherent with those observed in elderly or middle-aged human cohorts, and provide further information about earlier brain alterations and the pattern of disconnection associated with AD progression. Furthermore, the use of a rat model allows for better behavioral characterization than other animal models (Do Carmo & Cuello, 2013). This makes possible to perform cognitive evaluation of memory related functions and relate animal performance with brain connectivity. Our results revealed that the influence of brain network organization in cognitive abilities differs between transgenic and control animals.

Regarding the limitations of the study, the relatively small number of subjects could limit the statistical analysis. However, the measures were repeated at 5 time points, which increases the sample size evaluated by the longitudinal models to 80 observations. Note that our main results are based on models fitted to all these observations. Small sample could have a bigger impact on the complementary analysis at specific time points, but even with this limitation significant differences between the groups were detected after multiple comparison correction. These differences were coherent with previous findings observed in bigger cohorts from human population. Nevertheless, the small sample size could hamper the statistical analysis and be related, for instance, with the lack of significant differences at the last time point.

MRI protocols were optimized to achieve a compromise between sensitivity, image quality and acquisition time. After optimization, TE of the gradient-echo BOLD acquisition was set to 10.75 ms. Although it is slightly shorter than common echo-times used to sensitize images to BOLD variations, the analysis of the resulting BOLD signal showed patterns of connectivity consistent with previous knowledge, as shown in Supplementary Figure 2. Resting-state acquisitions were acquired using medetomidine as sedation, which was considered to preserve connectivity networks better than isoflurane anesthesia (Kalthoff, Po, Wiedermann, & Hoehn, 2013). However, recent studies have thoroughly investigated the effect of anesthesia in brain function during resting-state, and suggested that the use of only medetomidine could hinder brain function which has been observed in awake animals. These new findings should be taken into account in new experimental protocols, and could lead to additional conclusions that complement our analysis.

On the other hand, we would like to mention that all animals in the study repeated the DNMS task every three months, which allows for evaluation of cognitive skills. Results point to a learning effect and increase in cognitive reserve due to such repetition. However, further experiments including animals which do not perform DNMS should be carried out for a more thorough evaluation of such an effect.

Finally, the last time point evaluated in our study was 18 months of age. Brain changes and cognitive impairment in the TgF344-AD have been described mainly from 15 months of age, what, as previously discussed, could be related with the absence of differences in cognition or functional connectivity. Therefore, further investigation describing connectivity at later time points would be of great interest to characterize more advanced stages of the disease.

## CONCLUSIONS

Aging had more notable impact on the structural connectivity of the TgF344-AD rats, which is altered from early ages, than in control animals. Besides, differences in anatomical networks directly affected the cognitive outcome of the transgenic animals, even before the symptomatic onset. These findings are in line with results observed in middle-aged or elderly human population at risk of AD, and complement them with insights into earlier stages and a plot of the effects of the disease along the whole life span. The results support the idea of AD as a disconnection syndrome and AD as a continuum, suggesting that brain damage is already present at early stages, long before the symptomatic onset.

The impact of the altered anatomical connectivity in cognitive skills could be moderated by functional network reorganization until advanced stages of the disease. This suggests the relevance of cognitive reserve to prevent or mitigate the symptomatic onset in subjects affected by the disease. TgF344-AD model could therefore be a convenient model to perform translational research of the impact of cognitive interventions in AD.

## ACKNOWLEDGMENTS

This work has been funded by grants PI14/00595 and PI18/00893, integrated into Plan Nacional I+D+I and co-funded by ISCIII-Subdirección General de Evaluatión and European Regional Development Fund (ERDF). It was also funded by Fundació La Marató de TV3 (201441 10). Consorcio Centro de Investigatión Biomédica en Red (CIBER) de Bioingeniería, Biomateriales y Nanomedicina (CIBER-BBN) is an initiative financed by the Instituto de Salud Carlos III with assistance from the ERDF. With the support of Secretaria d’Universitats i Recerca del Departament d’Empresa i Coneixement de la Generalitat de Catalunya (AGAUR 2017 SGR 01003). The work was also funded by InMind project financed by the European Community (FP7-HEALTH-2011.2.2.1-2, n 278850). TgF344-AD rats were obtained through the InMind Consortium following kind donation by Dr. T. Town.

## AUTHOR CONTRIBUTIONS

All authors were involved in the conception and design of the project and manuscript writing and editing. GS and XLG was involved in data acquisition. XLG was in charge of the animal care and cognitive training and evaluation. GS, RT and EMM contribute to data interpretation. EMM performed the MRI processing and connectivity analysis and was a major contributor to the writing of the manuscript. All authors read and approved the final manuscript.

