## Supplementary Material for "Brain connectivity during Alzheimer’s disease progression and its cognitive impact in a transgenic rat model"

**Supplementary Materials**

**Index**

1. Network-based statistic results
2. Figures
3. Tables

### 1. Network-based statistic results

Networks altered in TgF344-AD animals resulting from NBS analysis involved the following regions:

- FD-w connectome at 8 months of age:
  - Right hemisphere: accumbens, superior colliculus, midline dorsal and ventromedial thalamus, temporal association cortex, medial geniculate and caudate putamen.
  - Left hemisphere: corpus callosum and cingulate cortex.
  - Both hemispheres: insular cortices.
- FA-w connectome at 15 months of age:
  - Right hemisphere: accumbens, frontal association cortex and cingulate cortex
  - Left hemisphere: corpus callosum, auditory cortex, anterior commissure, piriform cortex, visual cortex, posterior hippocampus, hippocampus subiculum, periaqueductal gray and superior colliculus.
  - Both hemispheres: entorhinal cortex, insular cortex, orbitofrontal cortex, caudate putamen, parietal association and somatosensory cortex, motor cortex, amygdala, IPAC and restrosplenial cortex.
- Functional connectome at 15 months of age:
  - Right hemisphere: midline dorsal and ventromedial thalamus, olfactory tubercle and visual cortex.
  - Left hemisphere: parietal association and somatosensory cortex.

**2. Figures**

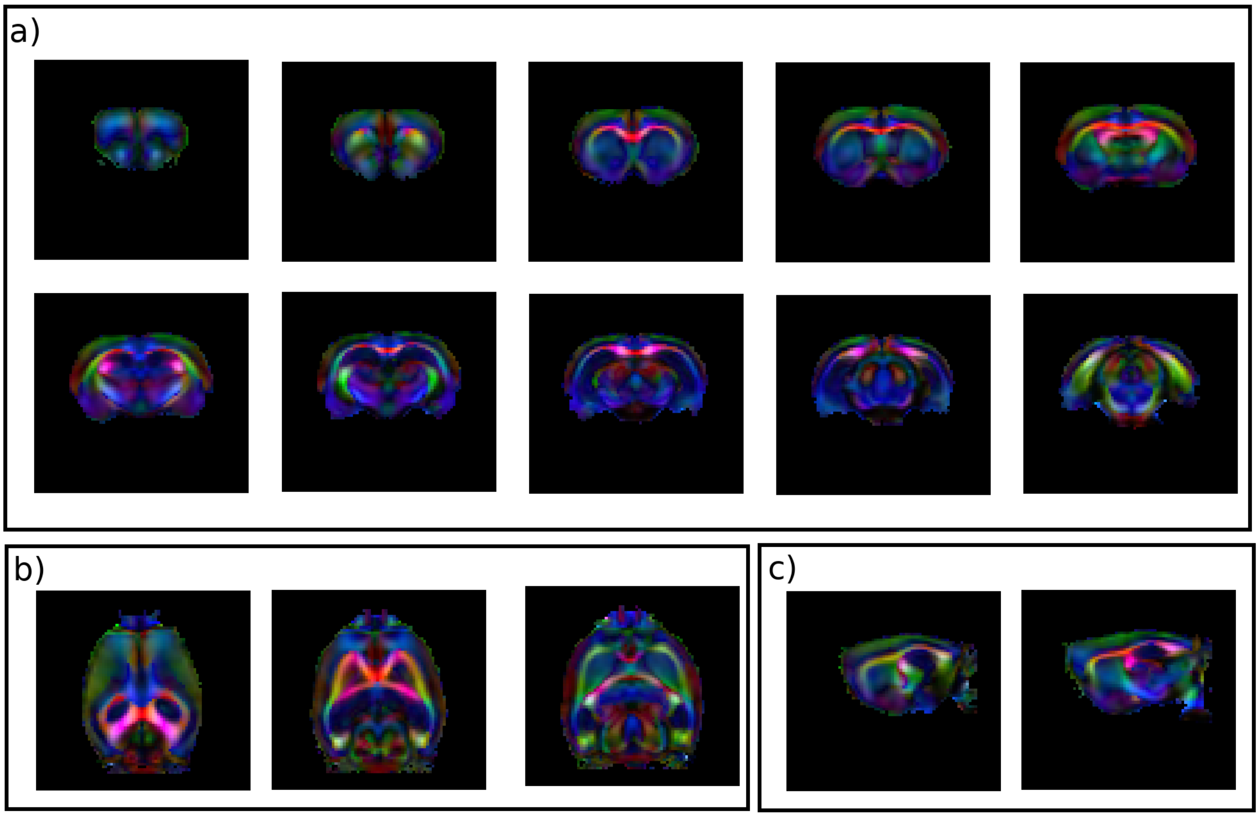


***Supplementary Figure 1.*** Representative slices of colored fractional anisotropy of one of the diffusion acquisitions. Color code: red, right-left direction; green, ventral-dorsal direction; blue: rostral-caudal direction. a) coronal slices from anterior to posterior; b) axial slices from superior to inferior; c) sagittal slices from left to right (left hemisphere).

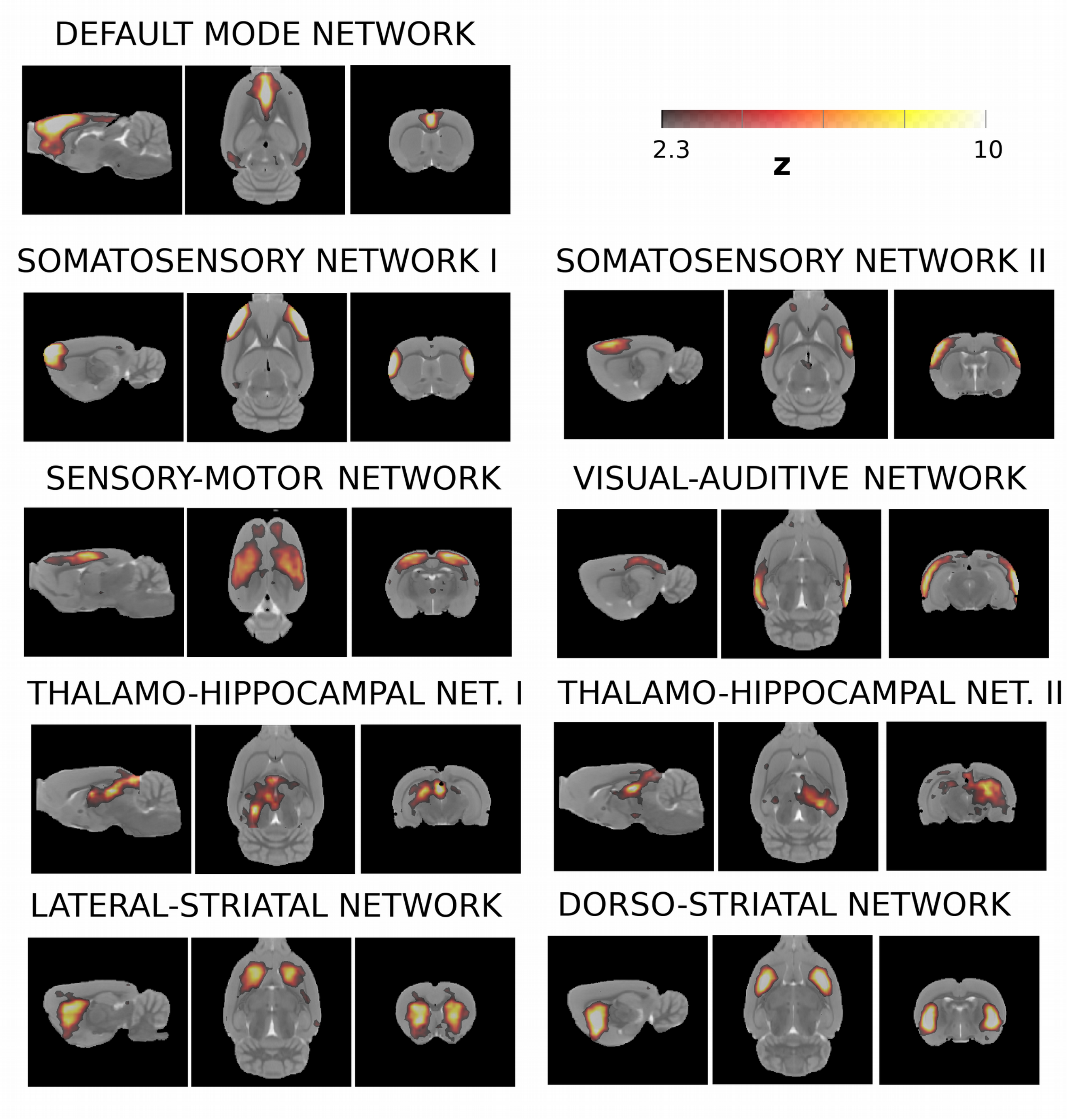


***Supplementary Figure 2.*** Representative slices of networks obtained by independent component analysis (ICA) of the dataset. ICA was performed using FLS's MELODIC (Jenkinson, Beckmann, Behrens, Woolrich, & Smith, 2012). The analysis was set to extract 30 components. The 9 networks shown in this figure were selected based on its correspondence to networks described in literature (Bajic, Craig, Borsook, Becerra, & Sullivan, 2016; Hsu et al., 2016; Sierakowiak et al., 2015).

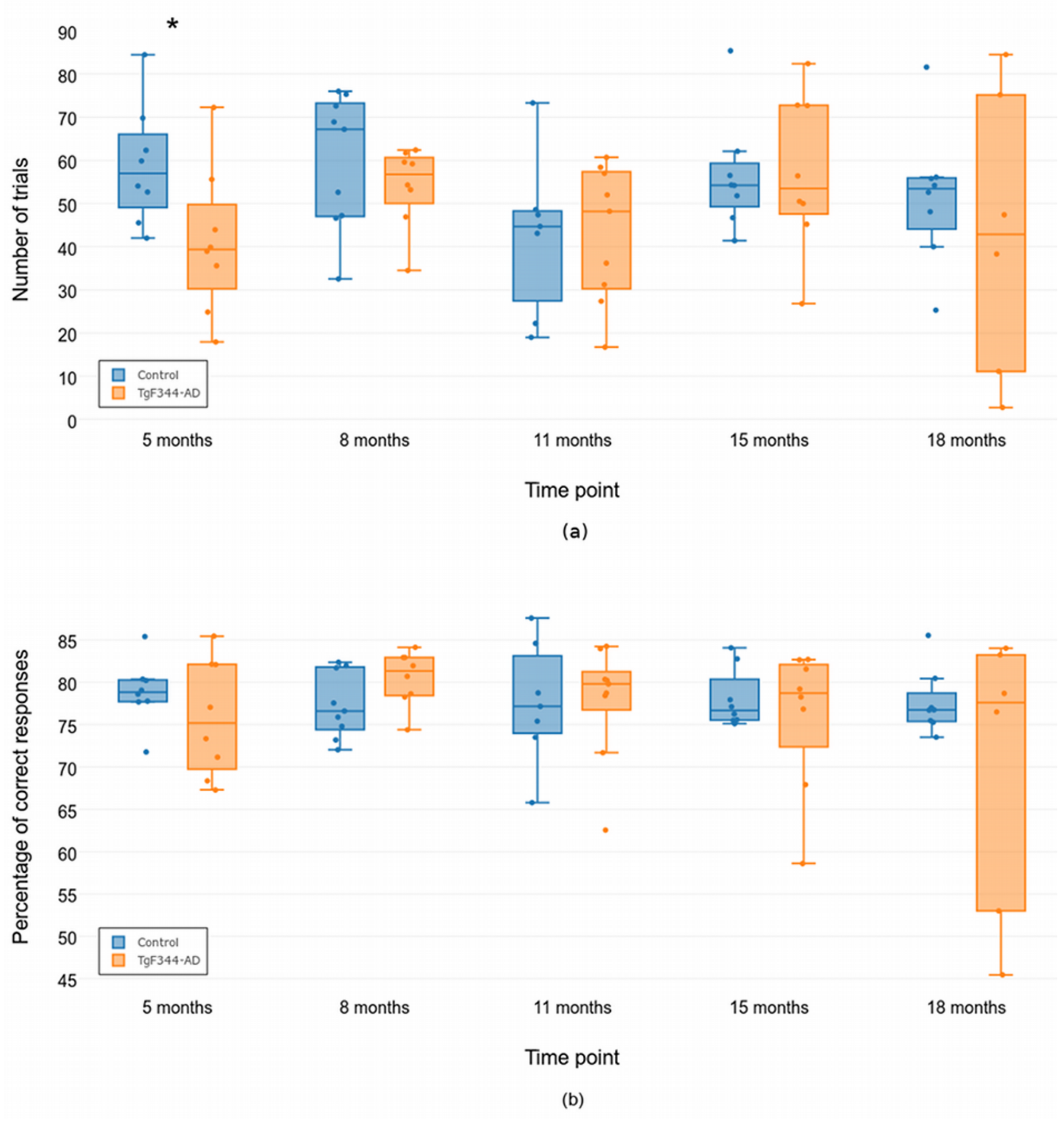


***Supplementary Figure 3.*** Cognitive performance. (a) Average number of trials per DNMS session and (b) percentage of correct responses in the DNMS task in the five evaluated time points (blue: control group; orange: TgF344-AD group). Asterisk represents uncorrected $p<0.05$. No significant differences were observed after FDR correction.

### 3. Tables

| | | Group | | Age | | Group-age interaction | |
|---|---|---|---|---|---|---|---|
| **Connectome** | **Network metric** | **$p_{FDR}$** | **Cohen's $f^2$** | **$p_{FDR}$** | **Cohen's $f^2$** | **$p_{FDR}$** | **Cohen's $f^2$** |
| FA-weighted | Strength | 0.0084* | 0.2553** | 0.0285* | 0.2143** | 0.0946 | 0.5226*** |
| | Global efficiency | 0.1164 | 0.2271** | 0.1149 | 0.1284* | 0.5259 | 0.1004* |
| | Local efficiency | 0.3012 | 0.1934** | 0.4528 | 0.0425* | 0.9151 | 0.0039 |
| | Clustering coeff. | 0.0266* | 0.2838** | 0.0400* | 0.2112** | 0.2772 | 0.2984** |
| FD-weighted | Strength | 0.1607 | 0.2029** | 0.1088 | 0.1209* | 0.0323* | 0.8118*** |
| | Global efficiency | 0.4155 | 0.1958** | 0.3323 | 0.0528* | 0.1378 | 0.4246*** |
| | Local efficiency | 0.2479 | 0.2135** | 0.1469 | 0.0961* | 0.0460* | 0.7060*** |
| | Clustering coeff. | 0.1164 | 0.1491* | 0.0673 | 0.1536** | 0.0276* | 0.9047*** |
| Structural binary | Degree | 0.0084* | 0.2305** | 0.0285* | 0.2383** | 0.0276* | 0.9597*** |
| | Global efficiency | 0.0084* | 0.2305** | 0.0285* | 0.2383** | 0.0276* | 0.9599*** |
| | Local efficiency | 0.0084* | 0.2186** | 0.0083* | 0.3353** | 0.0276* | 0.9154*** |
| | Clustering coeff. | 0.0084* | 0.2186** | 0.0083* | 0.3352** | 0.0276* | 0.9154*** |
| Functional weighted | Strength | 0.9056 | 0.0154 | 0.4716 | 0.0280* | 0.9151 | 0.0071 |
| | Global efficiency | 0.4262 | 0.0274* | 0.5612 | 0.0133 | 0.7252 | 0.0439* |
| | Local efficiency | 0.8754 | 0.0076 | 0.7547 | 0.0029 | 0.9744 | 0.0001 |
| | Clustering coeff. | 0.8754 | 0.0041 | 0.6528 | 0.0073 | 0.7932 | 0.0172 |
| Functional binary | Degree | 0.3012 | 0.0504 | 0.4708 | 0.0281* | 0.2772 | 0.2743** |
| | Global efficiency | 0.3012 | 0.0504 | 0.4708 | 0.0281* | 0.2772 | 0.2743** |
| | Local efficiency | 0.4155 | 0.0270 | 0.4935 | 0.0194 | 0.4530 | 0.1471* |
| | Clustering coefficient | 0.4155 | 0.0264 | 0.4935 | 0.0193 | 0.4530 | 0.1434* |

**Supplementary Table 1.** FDR corrected p-values and effect size quantified by Cohen's $f^2$ of the factors of the linear mixed effects (LME) model fitting each global network metric. In the $p_{FDR}$ columns * stands for statistical significance ($p_{FDR}<0.05$); in the Cohen's $f^2$ columns, asterisks represent small (*), medium (**) or large (***) effect size according to convention ($f^2 >0.02$, $f^2 >0.15$, $f^2 >0.35$, respectively).

| | | Control | | | TgF344-AD | | |
|---|---|---|---|---|---|---|---|
| **Connectome** | **Network metric** | **$p_{FDR}$** | **Cohen's $f^2$** | **$R^2$** | **$p_{FDR}$** | **Cohen's $f^2$** | **$R^2$** |
| FD-weighted | Strength | 0.4110 | 0.0734** | 0.0687 | 0.0462* | 0.1076** | 0.1328 |
| | Local efficiency | 0.4110 | 0.0695** | 0.0652 | 0.0803 | 0.0849** | 0.1547 |
| | Clustering coefficient | 0.4110 | 0.0655** | 0.0615 | 0.0462* | 0.1412** | 0.0873 |
| Structural binary | Degree | 0.4110 | 0.0345** | 0.1272 | 0.0462* | 0.0168 | 0.1440 |
| | Global efficiency | 0.4110 | 0.0345** | 0.1272 | 0.0462* | 0.1683** | 0.1440 |
| | Local efficiency | 0.7274 | 0.0051* | 0.1831 | 0.0420* | 0.2352** | 0.1904 |
| | Clustering coefficient | 0.7274 | 0.0051* | 0.1831 | 0.0420* | 0.2352** | 0.1904 |

**Supplementary Table 2.** Effect of age on the network metrics in each of the groups. FDR corrected p-values, effect size quantified by Cohen's $f^2$ and $R^2$ of the linear mixed effects (LME) model fitting each global network metric in control and transgenic animals independently. In the $p_{FDR}$ columns * stands for statistical significance ($p_{FDR}<0.05$); in the Cohen's $f^2$ columns, asterisks represent small (*), medium (**) or large (***) effect size according to convention ($f^2 >0.02$, $f^2 >0.15$, $f^2 >0.35$, respectively). Group models were evaluated only in the case of significant effect of the interaction between group and age in the LME model fitted to the whole cohort.

| | | Timepoint | | | | | | | | | |
|---|---|---|---|---|---|---|---|---|---|---|---|
| | | **1** (5 months) | | **2** (8 months) | | **3** (11 months) | | **4** (15 months) | | **5** (18 months) | |
| **Connectome** | **Network metric** | $p_{FDR}$ | $\eta^2$ | $p_{FDR}$ | $\eta^2$ | $p_{FDR}$ | $\eta^2$ | $p_{FDR}$ | $\eta^2$ | $p_{FDR}$ | $\eta^2$ |
| FA-weighted | Strength | 0.0371* | 0.5462*** | 0.1277 | 0.2889*** | 0.9578 | -0.0664 | 1.0000 | -0.0667 | 0.5474 | 0.0556* |
| | Global efficiency | 0.0566 | 0.3099*** | 0.2863 | 0.0722** | 0.9578 | -0.0616 | 0.2637 | 0.1333** | 0.5474 | 0.0556* |
| | Local efficiency | 0.1261 | 0.1303** | 0.2863 | 0.0377* | 0.9578 | -0.0616 | 0.0994 | 0.2889*** | 0.5474 | 0.0056 |
| | Clustering coefficient | 0.0371* | 0.4210*** | 0.1646 | 0.2056*** | 0.9578 | -0.0664 | 0.5802 | 0.0222* | 0.5474 | 0.0556* |
| FD-weighted | Strength | 0.1068 | 0.1838*** | 0.1277 | 0.2599*** | 0.9578 | -0.0696 | 0.0333* | 0.4525*** | 0.5474 | -0.0486 |
| | Global efficiency | 0.1261 | 0.1302** | 0.0542 | 0.5625*** | 0.9578 | -0.0696 | 0.1022 | 0.2599*** | 0.5474 | -0.0486 |
| | Local efficiency | 0.0971 | 0.2130*** | 0.1277 | 0.2889*** | 0.9578 | -0.0472 | 0.0333* | 0.4889*** | 0.5474 | -0.0486 |
| | Clustering coefficient | 0.0848 | 0.2437*** | 0.2863 | 0.1117** | 0.9578 | 0.0344* | 0.0333* | 0.4889*** | 0.6056 | -0.0611 |
| Structural binary | Degree | 0.0371* | 0.3824*** | 0.2863 | 0.0543* | 0.9578 | -0.0712 | 0.5802 | -0.0049 | 0.5474 | -0.0153 |
| | Global efficiency | 0.0371* | 0.3824*** | 0.2863 | 0.0543* | 0.9578 | -0.0712 | 0.5802 | -0.0049 | 0.5474 | -0.0153 |
| | Local efficiency | 0.0371* | 0.4210*** | 0.2863 | 0.0377* | 0.9578 | -0.0552 | 0.5802 | 0.0080 | 0.5474 | 0.0556* |
| | Clustering coefficient | 0.0371* | 0.4210*** | 0.2863 | 0.0377* | 0.9578 | -0.0552 | 0.5802 | 0.0080 | 0.5474 | 0.0556* |
| Functional weighted | Strength | 1.556 | 0.1380** | 0.3144 | 0.0222* | 0.9578 | -0.0472 | 1.0000 | -0.0642 | 0.5474 | 0.1167** |
| | Global efficiency | 0.1068 | 0.2488*** | 0.4589 | -0.0167 | 0.9578 | -0.0264 | 1.0000 | -0.0667 | 0.5474 | 0.0056 |
| | Local efficiency | 0.1989 | 0.0905** | 0.2863 | 0.0377* | 0.9578 | -0.0616 | 1.0000 | -0.0660 | 0.5474 | 0.0292* |
| | Clustering coefficient | 0.3254 | 0.0114* | 0.2863 | 0.0543* | 0.9578 | -0.0664 | 1.0000 | -0.0642 | 0.5474 | -0.0333 |
| Functional binary | Degree | 0.1156 | 0.2214** | 0.7000 | -0.0568 | 0.9578 | -0.0586 | 1.0000 | -0.0568 | 0.5474 | 0.0423* |
| | Global efficiency | 0.1156 | 0.2214** | 0.7000 | -0.0568 | 0.9578 | -0.0586 | 1.0000 | -0.0568 | 0.5474 | 0.0423* |
| | Local efficiency | 0.5698 | -0.0677 | 0.6655 | -0.0512 | 0.9578 | -0.0136 | 1.0000 | -0.0667 | 0.5474 | -0.0333 |
| | Clustering coefficient | 0.5698 | -0.0677 | 0.6655 | -0.0512 | 0.9578 | -0.0136 | 1.0000 | -0.0667 | 0.5474 | -0.0333 |

**Supplementary Table 3.** Comparison between network metrics in control and transgenic groups. FDR corrected p-values, effect size quantified by $\eta^2$ of the Kruskal-Wallis statistic. In the $p_{FDR}$ columns * stands for statistical significance ($p_{FDR}<0.05$); in the $\eta^2$ columns, asterisks represent small (*), medium (**) or large (***) effect size according to convention ($\eta^2 >0.01$, $\eta^2>0.06$, $\eta^2>0.14$, respectively).

| | | Group | | Metric | | Group x metric | |
|---|---|---|---|---|---|---|---|
| **Connectome** | **Network metric** | **$p_{FDR}$** | **Cohen's $f^2$** | **$p_{FDR}$** | **Cohen's $f^2$** | **$p_{FDR}$** | **Cohen's $f^2$** |
| FA-weighted | Strength | 0.0203* | 0.3774*** | 0.0006* | 0.5495*** | 0.0223* | 64.1640*** |
| | Global efficiency | 0.4643 | 0.0835* | 0.1778 | 0.1069* | 0.4909 | 23.1250*** |
| | Local efficiency | 0.7994 | 0.0210* | 0.8542 | 0.0029 | 0.7634 | 0.0409* |
| | Clustering coefficient | 0.1525 | 0.2020** | 0.0015 | 0.3170*** | 0.1622 | 55.3820*** |
| FD-weighted | Strength | 0.0098 | 0.3072** | 0.0007 | 0.5465*** | 0.0016 | 10.5991*** |
| | Global efficiency | 0.0783 | 0.1382* | 0.0035 | 0.2527** | 0.1261 | 5.0143*** |
| | Local efficiency | 0.0098 | 0.3007** | 0.0005 | 0.7049*** | 0.0167 | 15.0485*** |
| | Clustering coefficient | 0.0051 | 0.4168*** | 1.97e-5 | 0.9083*** | 0.0065 | 18.2783*** |
| Structural binary | Degree | 0.0051 | 0.6597*** | 1.6e-5 | 0.7300*** | 0.0065 | 140.7851*** |
| | Global efficiency | 0.0051 | 0.6597*** | 1.6e-5 | 0.7301*** | 0.0065 | 674.7025*** |
| | Local efficiency | 0.0098 | 0.7585*** | 4.4e-6 | 0.7568*** | 0.0064 | 2530.3059*** |
| | Clustering coefficient | 0.0098 | 0.7585*** | 4.4e-6 | 0.7568*** | 0.0064 | 565.0728*** |
| Functional weighted | Strength | 0.0098 | 0.1780** | 0.1097 | 0.1401 | 0.0150 | 56.0604*** |
| | Global efficiency | 0.0149 | 0.1518** | 0.0677 | 0.2013** | 0.018 | 43.9520*** |
| | Local efficiency | 0.0124 | 0.1607** | 0.0458* | 0.1765** | 0.0167 | 49.6405*** |
| | Clustering coefficient | 0.1242 | 0.1298* | 0.1097 | 0.1159* | 0.1417 | 38.0199*** |
| Binary weighted | Strength | 0.7363 | 0.0312* | 0.3502 | 0.0727* | 0.7107 | 0.0126 |
| | Global efficiency | 0.7363 | 0.0312* | 0.3502 | 0.0727* | 0.7107 | 0.1067* |
| | Local efficiency | 0.4036 | 0.0337* | 0.5046 | 0.0540* | 0.3740 | 29.4767*** |
| | Clustering coefficient | 0.4036 | 0.0336* | 0.5046 | 0.0536* | 0.3740 | 3.1351*** |

**Supplementary Table 4.** FDR corrected p-values and effect size quantified by Cohen's $f^2$ of the factors of the linear mixed effects (LME) model fitting the cognitive outcome (number of total trials) as a function of group and each global network metric. In the $p_{FDR}$ columns * stands for statistical significance ($p_{FDR}<0.05$); in the Cohen's $f^2$ columns, asterisk represent small (*), medium (**) or large (***) effect size according to convention ($f^2 >0.02$, $f^2 >0.15$, $f^2 >0.35$, respectively).

| | | Control | | | TgF344-AD | | |
|---|---|---|---|---|---|---|---|
| **Connectome** | **Network metric** | **$p_{FDR}$** | **Cohen's $f^2$** | **$R^2$** | **$p_{FDR}$** | **Cohen's $f^2$** | **$R^2$** |
| FA-weighted | Strength | 0.9976 | 0.0192 | 0.5818 | 0.0018 | 0.3979*** | 0.5230 |
| FD-weighted | Strength | 0.9976 | 0.0153 | 0.5885 | 0.0019 | 0.3739 | 0.5425 |
| | Local efficiency | 0.9976 | 0.0061 | 0.5849 | 0.0014 | 0.4182*** | 0.5545 |
| | Clustering coefficient | 0.9976 | 0.0080 | 0.5857 | 6.54e-5 | 0.6385*** | 0.6146 |
| Structural binary | Degree | 0.9752 | 0.0674* | 0.5929 | 5.29e-5 | 0.6968*** | 0.6257 |
| | Global efficiency | 0.9752 | 0.0674* | 0.5929 | 5.29e-5 | 0.6970*** | 0.6257 |
| | Local efficiency | 0.9752 | 0.0672* | 0.5953 | 1.26e-5 | 0.8208*** | 0.6508 |
| | Clustering coefficient | 0.9752 | 0.0672* | 0.5953 | 1.26e-5 | 0.8208*** | 0.6508 |
| Functional weighted | Strength | 0.0488 | 0.1753** | 0.5982 | 0.1570 | 0.1155* | 0.4139 |
| | Global efficiency | 0.1689 | 0.1107* | 0.5888 | 0.1088. | 0.1498* | 0.4383 |
| | Local efficiency | 0.1689 | 0.1110* | 0.5780 | 0.0771 | 0.1514** | 0.4254 |

**Supplementary Table 5.** Effect of network metrics in the cognitive outcome (number of total trials) of each group. FDR corrected p-values, effect size quantified by Cohen's $f^2$ and $R^2$ of the linear mixed effects (LME) model fitting cognitive outcome as function of network metrics in control and transgenic animals independently. In the $p_{FDR}$ columns * stands for statistical significance ($p_{FDR}$<0.05); in the Cohen's $f^2$ columns, asterisk represent small (*), medium (**) or large (***) effect size according to convention ($f^2$ >0.02, $f^2$ >0.15, $f^2$ >0.35, respectively). Group models were evaluated only in the case of significant effect of the interaction between group and network metric in the LME model fitted to the whole cohort.
